# Adding gene transcripts into genomic prediction improves accuracy and reveals sampling time dependence

**DOI:** 10.1101/2022.04.12.488053

**Authors:** B.C. Perez, M.C.A.M. Bink, K.L. Svenson, G.A. Churchill, M.P.L. Calus

## Abstract

Recent developments allowed generating multiple high quality ‘omics’ data that could increase predictive performance of genomic prediction for phenotypes and genetic merit in animals and plants. Here we have assessed the performance of parametric and non-parametric models that leverage transcriptomics in genomic prediction for 13 complex traits recorded in 478 animals from an outbred mouse population. Parametric models were implemented using best linear unbiased prediction (BLUP), while non-parametric models were implemented using the gradient boosting machine algorithm (GBM). We also propose a new model named GTCBLUP that aims to remove between-omics-layer covariance from predictors, whereas its counterpart GTBLUP does not do that. While GBM models captured more phenotypic variation, their predictive performance did not exceed the BLUP models for most traits. Models leveraging gene transcripts captured higher proportions of the phenotypic variance for almost all traits when these were measured closer to the moment of measuring gene transcripts in the liver. In most cases, the combination of layers was not able to outperform the best single-omics models to predict phenotypes. Using only gene transcripts, the GBM model was able to outperform BLUP for most traits except body weight, but the same pattern was not observed when using both SNP genotypes and gene transcripts. Although the GTCBLUP model was not able to produce the most accurate phenotypic predictions, it showed highest accuracies for breeding values for 9 out of 13 traits. We recommend using the GTBLUP model for prediction of phenotypes and using the GTCBLUP for prediction of breeding values.

## INTRODUCTION

Predicting complex traits is a fundamental aim of quantitative genetics. The use of whole genome single nucleotide polymorphisms (SNP) revolutionized the prediction of breeding values, resulting in the process widely known as genomic prediction (GP) (Meuwissen *et al*. 2001). A number of statistical approaches are now applied routinely in breeding programs, such as genomic best linear unbiased prediction (GBLUP) (VanRaden 2008), ridge regression (Whittaker *et al*. 2000), or methods from the “Bayesian Alphabet” (Gianola *et al*. 2009). More recently, machine learning algorithms have been tested in the context of genomic prediction (González-Recio *et al*. 2013; Pook *et al*. 2020; Zingaretti *et al*. 2020). These models may have several advantages when compared to traditional linear models, such as capturing interactions between predictors (non-additive effects), automatic variable selection and for making fewer assumptions regarding the underlying genetic architecture of phenotypes (Nayeri *et al*. 2019; Pérez-Enciso and Zingaretti 2019). However, compared to the linear models mentioned above, prediction performance from machine learning methods has shown mixed results (Azodi *et al*. 2019; Abdollahi-Arpanahi *et al*. 2020; Perez *et al*. 2022). There seems to be no “one-size-fits-all” model, as results are dependent on trait genetic architecture, size of the data, and on fine tuning of hyperparameters.

Recent development of low-cost high throughput molecular technologies allowed generating multiple high quality ‘omics’ data can be measured with high accuracy (Fernie and Schauer 2009; Tohge and Fernie 2015; Chawade *et al*. 2016). This has led to interest in utilizing these as new layers of information to improve the predictive performance of genomic prediction models, ultimately contributing to improve efficiency of breeding programs (Guo *et al*. 2016; Li *et al*. 2019). For example, gene expression levels measured in tissue samples by direct RNA sequencing (RNA-seq) is now readily available to animal breeders (Ozsolak and Milos 2011). To incorporate these new sources of data into genomic prediction models requires new strategies for integration with the already widely used genome-wide marker data. Although most of the literature focusing on the inclusion of gene-expression data into genomic models to improve predictive performance aimed at predicting phenotypes (Takagi *et al*. 2014; Guo *et al*. 2016; Schrag *et al*. 2018; Azodi *et al*. 2019; Li *et al*. 2019; Morgante *et al*. 2020), fitting gene transcript levels as an additional layer of information into genomic models could indirectly improve the prediction of breeding values. Christensen *et al*. (2021) presented a two-step method to incorporate such intermediate omics into genomic evaluations considering complete and incomplete omics-data scenarios. Results were validated using simulated data and suggested superiority of the single-step method including both the intermediate omics and genomics data, over the traditional genomic best linear unbiased prediction (GBLUP) using only genomics data. Similar results were observed by Michel *et al*. (2021) when investigating the integration of gene expression into genomic prediction for disease resistance in wheat by using a hybrid relationship matrix for merging both layers of omics data. A pending issue that remains, is the adequate handling of associations between layers of data that may lead to inflated relative contributions of individual layers when ignored (Holm *et al*. 2010; Christensen *et al*. 2021). Wade *et al*. (2021) have suggested that the benefits of multi-omics integration models over single-omic models are achieved once redundancy of predictors is decreased. Therefore, multi-omics models should either automatically or through adequate parametrization be able to identify and manage information redundancy across multiple omics-layers.

In the present study we used data from the Diversity Outbred (DO) mouse population (Churchill *et al*. 2012; Svenson *et al*. 2012) to evaluate the utility of gene expression in addition to genome-wide genetic markers for genomic prediction using different modeling strategies. To this end, the objectives of this study were to: (1) assess the proportions of phenotypic variance explained by genetic markers and gene transcripts in complex traits recorded in at least two time points; (2) evaluate the predictive accuracy for phenotypes using transcripts and/or marker information for the traits investigated using linear models and the gradient boosting algorithm; and (3) evaluate how the inclusion of transcripts affects estimation of genomic breeding values (GEBV) from BLUP models. The linear models proposed vary in number of components, how interactions were modeled, and conditioning of one component on another. The gradient boosting machine algorithm was chosen for its ability to automatically control redundancy and implicitly account for non-linear effects in prediction, while the BLUP models tested comprise parametric approaches to incorporate genomics and transcriptomics, considering or ignoring the interactions between them.

## MATERIAL AND METHODS

### Data

#### Phenotypes

Data used for this study were obtained from The Jackson Laboratory (Bar Harbor, ME) and comprise a subset of the dataset used in Perez *et al*. (2022). The 478 DO mice originated from 4 non-overlapping generations (4, 5, 7 and 11) with males and females represented equally. The total number of animals per generation was 47, 47, 192 and 192 for generations 4, 5, 7 and 11, respectively, with slight variation in the numbers of missing records across traits (Table 1). The mice were maintained on either standard high fiber (chow, n=239) or high fat diet (n=239) from weaning until 23 weeks of age. The proportion of males and females within each diet category was close to 50-50 for all generations, as well as within each litter-generation combination (two litters per generation). This population is maintained under a systematic mating scheme, designed to limit population structure and relatedness. On average, the animals were related to each other at a level equivalent to first cousins, which is by design (Svenson *et al*. 2012). More elaborate descriptions of population structure, husbandry and phenotyping methods can be found in Svenson *et al*. (2012) and Tyler *et al*. (2021).

**Table 1.** Number of available observations (N), the extended description of traits, age of the animals at phenotype measurement, and estimated heritability.

| Trait | N | Trait description | Age at measurement | Estimated heritability<sup>1</sup> |
| --- | --- | --- | --- | --- |
| BMD12 | 471 | Bone mineral density | 12 weeks | 0.39 |
| BMD21 | 471 | Bone mineral density | 21 weeks | 0.41 |
| BW10 | 478 | Body weight | 10 weeks | 0.42 |
| BW15 | 478 | Body weight | 15 weeks | 0.35 |
| BW20 | 478 | Body weight | 20 weeks | 0.37 |
| CHOL8 | 474 | Circulating cholesterol | 8 weeks | 0.38 |
| CHOL19 | 474 | Circulating cholesterol | 19 weeks | 0.45 |
| FATP12 | 471 | Body fat percentage | 12 weeks | 0.35 |
| FATP21 | 471 | Body fat percentage | 21 weeks | 0.32 |
| GLUC8 | 425 | Circulating glucose | 8 weeks | 0.31 |
| GLUC19 | 425 | Circulating glucose | 19 weeks | 0.22 |
| TRGL8 | 473 | Circulating triglycerides | 8 weeks | 0.36 |
| TRGL19 | 473 | Circulating triglycerides | 19 weeks | 0.31 |

<sup>1</sup> Standard errors for the heritability ranged from 0.07 to 0.09

Table 1 gives for each trait a brief description, the numbers of observations and the estimated heritability. We considered six traits based on range of heritability and presumed genetic architectures. The chosen traits were measured at two or three times during the animal’s life, resulting in 13 distinct traits in total. The analyzed traits were bone mineral density at 12 (BMD12) and 21 (BMD21) weeks, body weight at 10, 15 and 20 weeks (BW10, BW15 and BW20); circulating cholesterol at 8 (CHOL8) and 19 (CHOL19) weeks, adjusted body fat percentage at 12 (FATP12) and 21 (FATP21), circulating glucose at 8 (GLUC8) and 19 (GLUC19) weeks, circulating triglycerides at 8 (TRGL8) and 19 weeks (TRGL19). These traits can be categorized into measurements of body composition (bone mineral density, body weights and fat percentage) and clinical plasma chemistries (circulating glucose and triglycerides). Phenotypic records were pre-corrected for fixed effects of diet, generation, litter, and sex (Perez *et al*. 2022). Therefore, the pre-corrected phenotypes (**y**^*^) analyzed here comprise the sum of the additive genetic effect and residual terms.

#### Genotypes

The genotype data used for the animals in this study, were obtained from their derived founder haplotypes (for details: see Perez *et al*. 2022). The complete genotype file used for the analyses included 64,000 markers on an evenly spaced grid, and the average distance between markers was 0.0238 cM. The full genotype dataset was cleaned based on the following criteria: variants with minor allele frequency < 0.05, call rates < 0.90 and linear correlation between subsequent SNPs > 0.80 were removed. After quality control, a total of 50,122 SNP markers were available for the mice with phenotypic, genotypic and transcriptomic records.

#### Transcript levels

Transcriptome-wide expression levels were measured from whole livers as previously described (Munger *et al*. 2014; Chick *et al*. 2016) for 478 animals at 26 weeks of age. The RNA sample was sequenced using single-end RNA-Seq (Munger *et al*. 2014) and aligned transcripts to strain-specific genomes from the DO founders (Chick *et al*. 2016). Read counts were estimated using an expectation maximization algorithm (EMASE, https://github.com/churchill-lab/emase). Read counts were previously corrected for the effects of sex, diet, and batch by normalizing in each sample using upper quantile normalization and applying a rank Z transformation across samples. After quality control, quantification of transcripts was available for 11,770 genes (Tyler *et al*. 2017).

#### Statistical models

Below we introduce five best linear unbiased prediction (BLUP) models and three gradient boosting machine (GBM) models with their acronyms and key features summarized in Table 2.

**Table 2.**
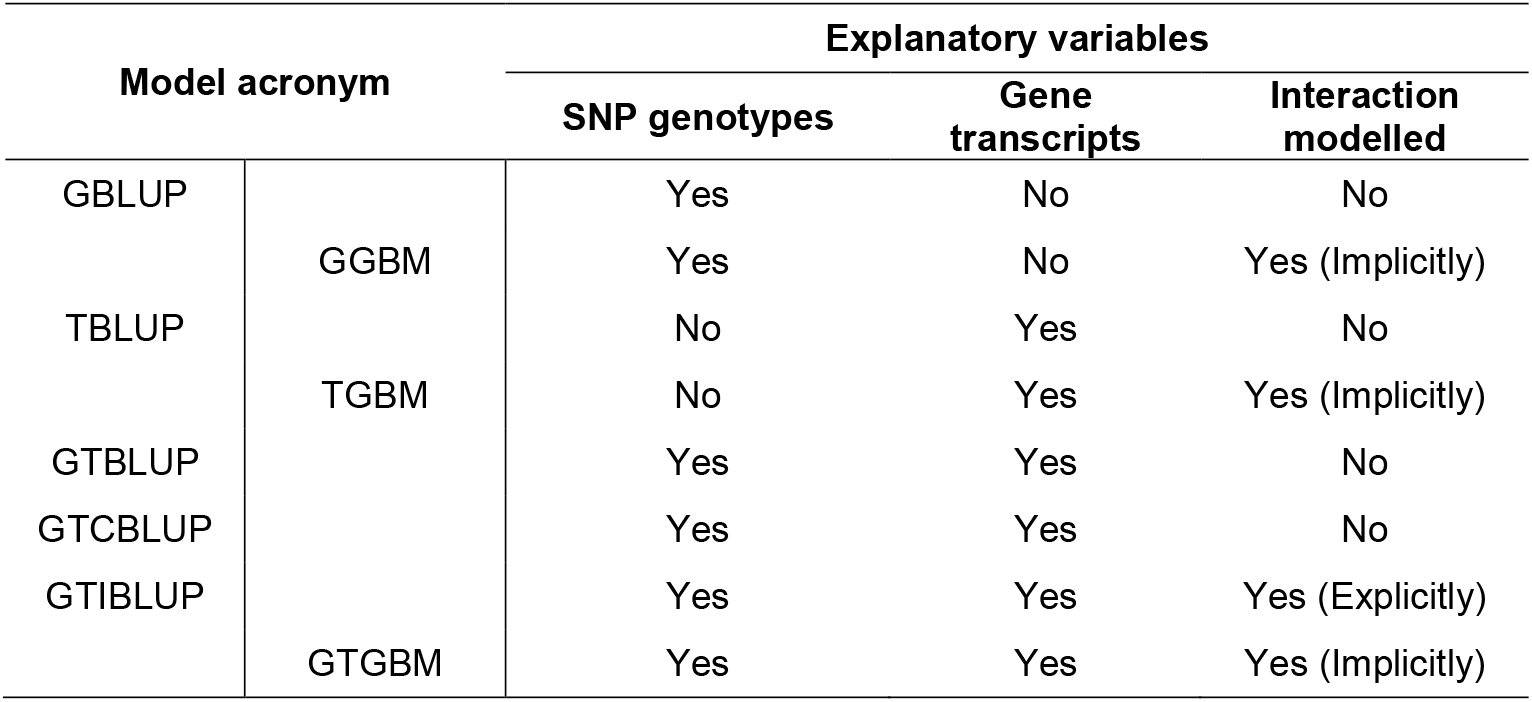
Overview of models applied to SNP genotypes and/or individual levels of gene transcripts.

#### Best linear unbiased prediction

### GBLUP

The statistical model of GBLUP is:

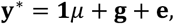

where **y**^*^ is the vector of pre-corrected phenotypes, **1** is a vector of ones, *μ* is the intercept, **g** is the vector of random additive genetic values, where 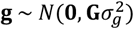, **G** is the additive genomic relationship matrix between genotyped individuals, and 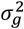 is the additive genomic variance. The matrix **G** is constructed following the second method described by VanRaden (2008) as 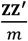 where **Z** is the matrix of centered and standardized genotypes for all individuals and *m* is the number of markers. Finally, **e** is the vector of random residual effects where 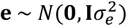 with 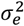 being the residual variance, and **I** is an identity matrix.

#### TBLUP

To evaluate the performance of transcriptomic data for predicting complex traits, we used a Transcriptomic Best Linear Unbiased Predictor (TBLUP) model. This model is similar to GBLUP, but using a transcriptomic relationship matrix, which evaluates the similarity among animals based on gene expression levels (Guo *et al*. 2016).

The statistical model of TBLUP is:

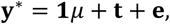

where **y**^*^, **1** and *μ* are defined as above, **t** is the vector of random transcript level effects, where 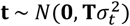 and **T** is the transcriptomic relationship matrix built according to the formula 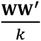 where **W** is the matrix of centered and standardized expression levels for all animals and *k* is the number of genes, and 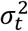 is the variance explained by gene transcripts.

#### GTBLUP and GTIBLUP

The GTBLUP model fitted the **g** and **t** as independent random effects, each with their own variance component (Guo *et al*. 2016; Li *et al*. 2019). The model is **y**^*^ = **1***μ* + **g** + **t** + **e**, where all the parameters are as defined above.

The GTIBLUP model fitted **g, t**, and the interaction between **g** and **t** with an additional variance component (Morgante *et al*. 2020). This model is **y**^*^ = **1***μ* + **g** + **t** + **gt** + **e**, where **y**^*^, **1***μ*, **g, t** and **e** are as defined above, and **gt** is the vector of interaction (between genomic and transcriptomic) effects, where 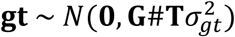 and # is the Hadamard product.

#### GTCBLUP

The GTCBLUP model was similar to GTBLUP in that the **g** and **t** that were fitted as independent random effects, each with their own variance component. However, for this model the transcript levels were conditioned on SNP genotypes, yielding a matrix **W**_***c***_ computed as: **W**_***c***_ = **(I** − **Z**(**Z**^′^**Z** + **I**λ)^−**1**^**Z**′)**W**, where **Z**(**Z**^′^**Z** + **I**λ)^−**1**^**Z**′ is the so-called “smoother matrix” (Hastie *et al*. 2009), **Z** is the matrix of centered and standardized genotypes as before, **I** is an identity matrix, and 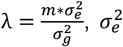 is the residual variance, and 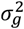 is the additive genomic variance, both variances estimated with the GBLUP model (including only **g**). Using the smoother matrix, i.e. including **I**λ rather than using **I** − **Z**(**Z**^′^**Z**)^−**1**^**Z**′, reflects that the effects associated with the SNPs are estimated as random rather than fixed effects. The aim of this model is to remove any variance from transcripts that is correlated to variance in genotypes, such that any phenotypic variance both associated with variance in genotypes and transcripts automatically will be associated with the genotypes only. The model is **y**^*^ = **1***μ* + **g** + **t**_***c***_ + **e**, where 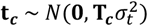 and **T**_***c***_ is computed as 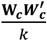, and all other parameters are as defined above.

#### Gradient boosting machine models

Gradient boosting machine (GBM) is an ensemble learning technique that applies an iterative process of assembling “weak learners” into a stronger learner, being largely used for both classification and regression problems (Friedman 2001). In the scope of this investigation, the GBM algorithm represents a non-parametric approach capable of implicitly fitting not only the additive effects of SNP and gene transcripts, but also the within- and between-omics layers interactions. The GBM is also capable of performing automatic feature selection, prioritization of important variables and discarding variables containing irrelevant or redundant information. A detailed description of the gradient boosting machine algorithm and its application in genomic prediction can be found in Friedman (2002), González-Recio *et al*. (2010; 2013) and Perez *et al*., (2022).

To obtain the best possible results from the GBM algorithm, a grid search approach was used to determine the combination of hyperparameters that minimized the mean squared error of prediction within the inner training set for each trait. Details of the hyperparameter search method used are found in Perez *et al*. (2022). We implemented the GBM model using the “gbm” R package (Ridgeway 2020).

We tested three different GBM models. The first model considered only SNP genotypes as predictors (GGBM), the second model considered only (standardized) gene transcript levels as predictors (TGBM) and a third model that considered both genetic markers and transcript levels together as predictors (GTGBM). Our objective was to investigate if GBM models could capture within and between omics layers associations, while also reducing within and between omics layers redundancy by performing automatic variable selection. It is important to note here that although here we used “G” and “T” letters to refer to genomics and transcriptomics data in the GBM model’s acronyms, predictors were fit directly in the model and not as relationship matrices

#### Variance explained by genetic markers, transcript levels and combinations of both

To understand how much of the phenotypic variance can be explained by using SNP genotypes, gene transcript levels and the combinations of both sources of information, we estimated variance components using the GBLUP, TBLUP, GTBLUP, GTIBLUP and GTCBLUP models. Estimates of variance components along with the residual variance 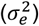 were obtained from a Bayesian approach analysis, using the BGLR R package (Pérez and de los Campos 2014). For all models, the Gibbs sampler was run for 60,000 iterations, with a 20,000 burn-in period and a thinning interval of 10 iterations. Consequently, inference was based on 4,000 posterior samples.

For the GTIBLUP model, we calculated the portion of variance explained by SNP genotypes 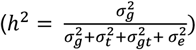, gene transcripts 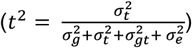 and from the interaction between effects from genetic markers and gene transcripts 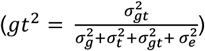 Consequently, the sum of *h*^2^, *t*^2^ and *gt*^2^ represent the portion of phenotypic variance explained by two layers of omics data and by the between-omics-layer interactions. The parameters *h*^2^, *t*^2^ and *gt*^2^ for the other models were calculated similarly but omitted any variance components associated with effects not included in the model.

#### Model Performance

Performance of predictions from the models was measured by the accuracy, computed as the Pearson correlation (r_**y**_^*^ _ŷ_), and the relative root-mean squared error of prediction (RRMSE) between predictions (ŷ) and pre-corrected phenotypes 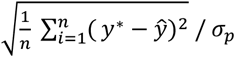, where σ_*p*_ is the phenotypic standard deviation. In all analyses, we used a forward prediction validation scheme in which animals from older generations (4, 5, 7) were used as the reference and animals from the younger generation (11) as the validation subset. The standard error (SE) around the r_**y**_^*^ _ŷ_ estimates were obtained by calculating the standard deviations from 10,000 bootstrap samples (Davison and Hinkley 1997). The bootstrapping procedure was implemented using the “boot” R package (Canty and Ripley 2021). We have also assessed prediction bias by obtaining the regression coefficient from the linear regression of corrected phenotypes on model predictions. For these results, values above 1 indicate deflation, while values below 1 indicate inflation of predicted values.

To assess the proportion of variance explained by the models tested, we have calculated the coefficient of determination (R^2^) from the regression of corrected phenotypes on model predictions for all traits. For the GBM models we have used results from the model using the previously obtained best hyperparameter set from the standard grid-search procedure to assess the model R^2^ for prediction within the reference set.

For the BLUP models proposed to integrate SNP genotypes and gene transcripts (GTBLUP, GTIBLUP and GTCBLUP), in addition to r_**y**_^*^, _ŷ_ we have also calculated the correlations between the solutions for each random effect included in the model (**g, t, t**_**c**_ or **gt**) and *ŷ*, as well as pairwise comparisons between all components in the model. Here, we also focus on solutions from the additive genetic component from these models to assess if the prediction of genomic breeding values (GEBV) can be improved by using models capable of integrating SNP genotypes and gene transcripts for genomic prediction.

### Data Availability

All data associated with this manuscript, and the code developed and used to perform analyzes described in this manuscript, can be obtained at https://doi.org/10.6084/m9.figshare.15081636.v1. All software used is publicly available.

## RESULTS

### Variance components estimation percentage of variance explained within the reference set

Genomic heritabilities (*h*^2^) obtained with GBLUP ranged from 0.08 to 0.44, representing a wide range of magnitudes across traits (Figure 1 and Figure 2). When only fitting transcript levels as predictors (TBLUP), the percentage of variance explained (*t*^2^) ranged from 0.22 to 0.75 and in general it was higher than *h*^2^ when comparing within the same trait. The exceptions to that were observed for BMD12 (*h*^2^ = 0.39 and *t*^2^ = 0.35) and GLUC8 (*h*^2^ = 0.30 and *t*^2^ = 0.22). When comparing the same trait measured at different time points, *t*^2^ from TBLUP was higher for phenotypes collected closer to the 26 weeks of age (i.e. the age at mRNA data sampling).

**Figure 1.**
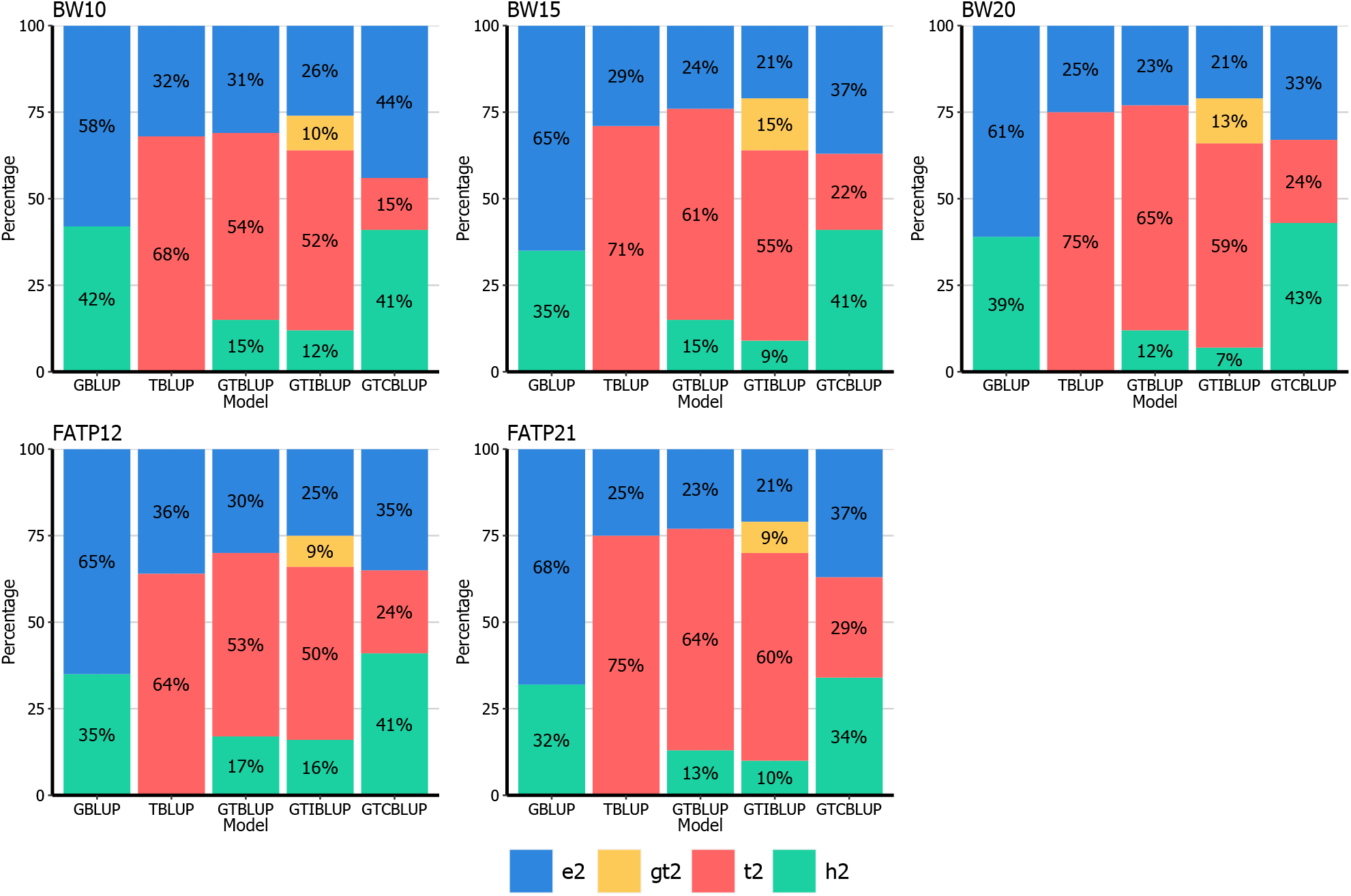
Percentage of variance explained by SNP genotypes (*g*^2^), gene transcripts (*t*^2^), the interaction between them (*gt*^2^) and not explained (*e*^2^) by GBLUP, TBLUP, GTBLUP, GTIBLUP and GTCBLUP models tested for the traits BW and FATP. ^1^For a description of the traits, see Table 1. ^2^For a description of the models, see Table 2.

**Figure 2.**
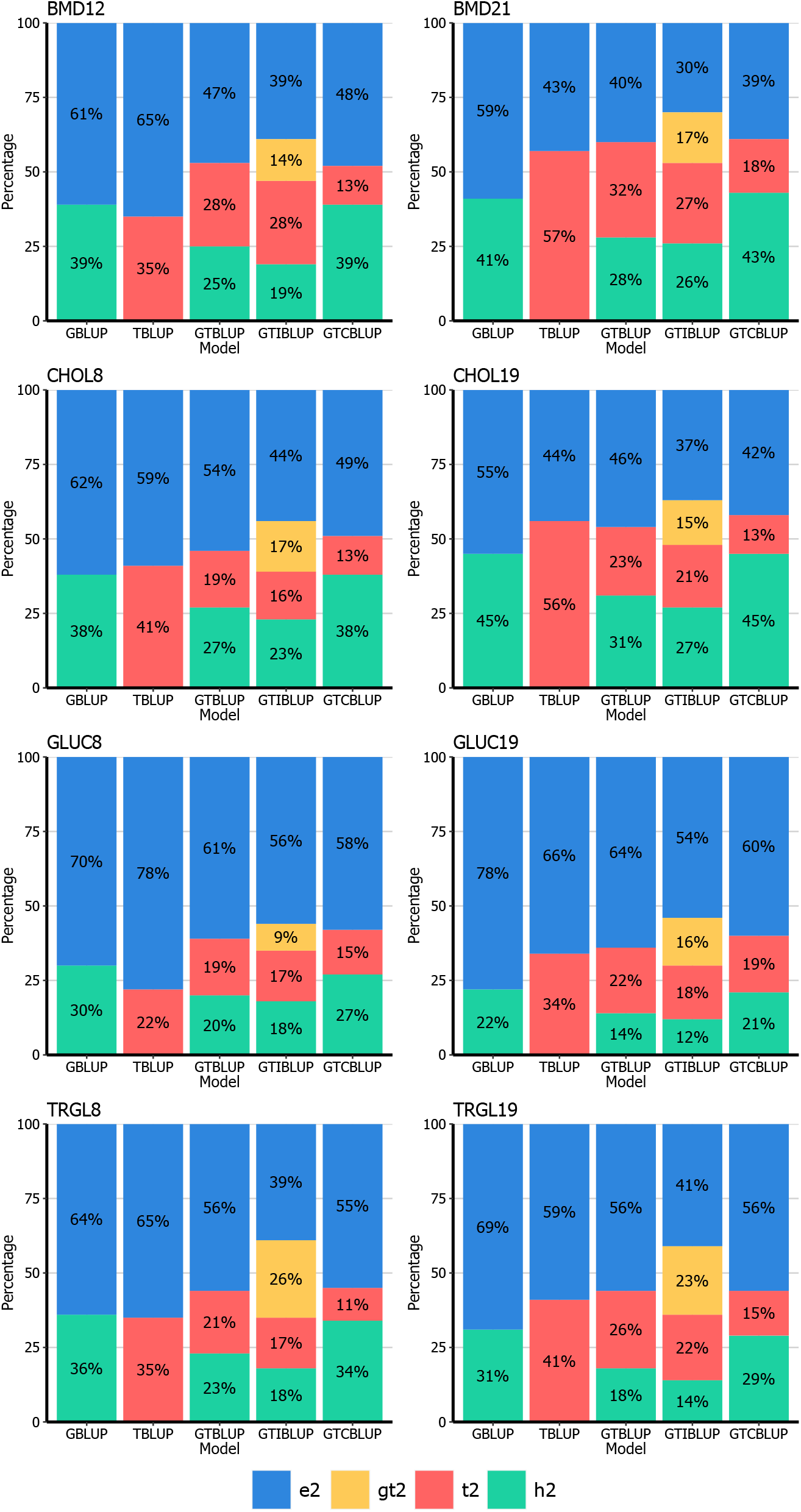
Percentage of variance explained by SNP genotypes (*g*^2^), gene transcripts (*t*^2^), the interaction between them (*gt*^2^) and not explained (*e*^2^) by GBLUP, TBLUP, GTBLUP, GTIBLUP and GTCBLUP models tested for the traits BMD, CHOL, GLUC and TRGL. ^1^For a description of the traits, see Table 1. ^2^For a description of the models, see Table 2.

In terms of the total phenotypic variance explained, GTBLUP and GTIBLUP showed similar results (Figure 1 and Figure 2). For body weights (BW10, BW15, BW20) and fat percentage (FATP12 and FATP21) traits, the variance explained by genetic markers (in GTBLUP and GTIBLUP) was drasticly lower when compared to GBLUP for the same traits. For the remaining traits the decrease in genetic variance captured by markers was much lower. For the interaction component in GTIBLUP (*gt*^2^), results observed varied according to the trait analysed but in general, it was low compared to *h*^2^ and *t*^2^. The only exception to that was observed for TRGL8, in which *gt*^2^ was higher than *h*^2^ and *t*^2^. For CHOL8, GLUC19 and TRGL19, *gt*^2^ was either similar to *h*^2^ or *t*^2^.

For GTCBLUP, differently from GTBLUP and GTIBLUP, the additive genetic variance captured was always in line with results from GBLUP. On the other hand, the variance explained by transcripts (*t*^2^) from GTCBLUP was always lower than observed by other models including transcripts as predictors (TBLUP, GTBLUP and GTIBLUP).

The variance explained (represented by the R^2^ parameter) within the reference data by parametric models was in general lower than by the non-parametric counterparts (Table 3). Independent of being a parametric or non-parametric model, the use of gene transcripts (TBLUP and TGBM) as predictors explained in most cases more of the variance than using exclusively SNP genotypes (GBLUP and GGBM). For GTBLUP, GTIBLUP and GTGBM, the variance explained was at least similar to observed for TBLUP and TGBM, but generally higher. For GTCBLUP, variance explained by the model was slightly to moderately higher than observed for GBLUP model, but always smaller than observed for GTBLUP, GTIBLUP and GTGBM. The average R^2^ when considering only traits recorded earlier (suffixes 8, 10 or 12) and later (suffixes 19, 20 or 21) moments were 76% and 83%, respectively, when using TBLUP, being the largest differece observed across models when considering these two groups of traits.

**Table 3.** Model R^2^ (x100) for the best linear unbiased prediction (GBLUP, GTBLUP, GTCBLUP and GTIBLUP) and gradient boosting (GGBM, TGBM and GTGBM) approaches within training data^1^.

| Trait<sup>1</sup> | Model<sup>2</sup> |  |  |  |  |  |  |  |
| --- | --- | --- | --- | --- | --- | --- | --- | --- |
|  | Only SNP |  | Only gene transcripts |  | SNP + gene transcripts |  |  |  |
|  | GBLUP | GGBM | TBLUP | TGBM | GTBLUP | GTIBLUP | GTCBLUP | GTGBM |
| BMD12 | 75 | 90 | 79 | 96 | 85 | 92 | 88 | 95 |
| BMD21 | 84 | 85 | 88 | 93 | 87 | 96 | 92 | 98 |
| BW10 | 78 | 92 | 93 | 96 | 91 | 95 | 90 | 97 |
| BW15 | 75 | 87 | 91 | 94 | 93 | 96 | 94 | 96 |
| BW20 | 80 | 87 | 92 | 93 | 95 | 97 | 95 | 97 |
| CHOL8 | 84 | 85 | 69 | 97 | 80 | 93 | 87 | 98 |
| CHOL19 | 82 | 83 | 85 | 95 | 88 | 96 | 92 | 98 |
| FATP12 | 75 | 76 | 89 | 97 | 92 | 96 | 95 | 98 |
| FATP21 | 80 | 82 | 93 | 97 | 95 | 97 | 94 | 98 |
| GLUC8 | 77 | 85 | 66 | 95 | 81 | 93 | 87 | 97 |
| GLUC19 | 62 | 80 | 67 | 96 | 75 | 90 | 84 | 98 |
| TRGL8 | 65 | 81 | 62 | 92 | 80 | 96 | 88 | 97 |
| TRGL19 | 71 | 86 | 70 | 96 | 78 | 95 | 84 | 98 |
| Mean (all) | 76 | 85 | 80 | 95 | 86 | 95 | 90 | 97 |
| Mean (T1) | 75 | 85 | 76 | 96 | 85 | 94 | 89 | 97 |
| Mean (T2) | 76 | 84 | 83 | 95 | 86 | 95 | 90 | 98 |

T1 = average R<sup>2</sup> of the column considering only traits recorded earlier in life (suffixes 8, 10 and 12).
T2 = average R<sup>2</sup> of the column considering only traits recorded later in life (suffixes 19, 20 and 21).
<sup>1</sup>For a description of the traits, see Table 1.
<sup>2</sup>For a description of the models, see Table 2.
<sup>3</sup>BW15 trait was ignored when calculating average performance considering exclusively T1 and T2.

#### Prediction performance – Phenotype prediction

In Table 4 accuracies are shown for predicted phenotypes for BLUP and GBM models using either SNP genotypes, transcript levels or both as predictors. Here we considered GBLUP to be the reference method. It showed prediction accuracies ranging from 0.01 to 0.29, these were highly positively correlated to the portion of variance explained by SNP genotypes by the same model, except for CHOL19. When comparing predictive performance between GBLUP and GGBM models, GBLUP yielded highest prediction accuracies for 7 traits, while GGBM had best predictive performance for 6 traits out of 13.

**Table 4.** Accuracies of predicted pre-corrected phenotypes for the validation subset with the proposed models. For each group of models, the result with the highest accuracy is indicated in bold, identical results between two or more models are indicated in underline.

| Trait<sup>1</sup> | Model<sup>2</sup> |  |  |  |  |  |  |  |
| --- | --- | --- | --- | --- | --- | --- | --- | --- |
|  | Only SNP |  | Only gene transcripts |  | SNP + gene transcripts |  |  |  |
|  | GBLUP | GGBM | TBLUP | TGBM | GTBLUP | GTIBLUP | GTCBLUP | GTGBM |
| BMD12 | **0.19** | 0.16 | **0.27** | 0.25 | <u>0.27</u> | <u>0.27</u> | 0.20 | 0.25 |
| BMD21 | **0.29** | 0.28 | <u>0.38</u> | <u>0.38</u> | **0.42** | 0.40 | 0.29 | 0.38 |
| BW10 | 0.18 | **0.20** | **0.48** | 0.42 | <u>0.47</u> | <u>0.47</u> | 0.19 | 0.42 |
| BW15 | **0.15** | 0.12 | <u>0.52</u> | <u>0.52</u> | <u>0.51</u> | <u>0.51</u> | 0.22 | 0.49 |
| BW20 | **0.18** | 0.16 | **0.61** | 0.58 | <u>0.60</u> | <u>0.60</u> | 0.30 | 0.54 |
| CHOL8 | 0.14 | **0.17** | 0.14 | **0.15** | **0.18** | 0.17 | 0.16 | 0.16 |
| CHOL19 | **0.25** | 0.20 | **0.19** | 0.16 | **0.26** | 0.23 | 0.22 | 0.23 |
| FATP12 | 0.14 | **0.15** | 0.44 | **0.45** | 0.44 | 0.44 | 0.28 | **0.46** |
| FATP21 | **0.21** | 0.20 | 0.54 | **0.56** | **0.54** | 0.53 | 0.35 | 0.52 |
| GLUC8 | 0.10 | **0.12** | 0.03 | **0.04** | 0.08 | 0.09 | 0.11 | **0.15** |
| GLUC19 | 0.01 | **0.10** | 0.03 | **0.05** | 0.04 | 0.05 | -0.05 | **0.11** |
| TRGL8 | 0.08 | **0.11** | 0.06 | **0.08** | **0.07** | 0.06 | 0.05 | 0.06 |
| TRGL19 | **0.15** | 0.13 | 0.17 | **0.19** | 0.17 | 0.18 | 0.12 | **0.19** |

<sup>1</sup>For a description of the traits, see Table 1.
<sup>2</sup>For a description of the models, see Table 2.

For models that include only gene transcripts (TBLUP and TGBM), the TBLUP approach showed predictive accuracies ranging from 0.03 to 0.61, having the best performance for only 4 out of 13 traits. The TGBM model was able to overcome TBLUP for 7 traits, with prediction accuracies ranging from 0.04 to 0.58. For BMD21 and BW15, predictive accuracy was identical between TBLUP and TGBM. The differences between accuracies from TBLUP and TGBM was higher than between GBLUP and GGBM.

For models that combined SNP genotypes and gene transcripts levels (GTBLUP, GTCBLUP, GTIBLUP and GTGBM), GTBLUP had the highest predictive accuracy for 5 traits out of 13. The second-best model overall was GTGBM, with the highest predictive accuracy for 4 traits. For every trait that GTIBLUP had the highest prediction accuracy, it was identical to the result for GTBLUP, while the GTCBLUP never had the highest predictive accuracy (Table 4).

The prediction error (RRMSE) and bias (*β*) for model’s predictions are presented in Supplementary Tables S1 and S2, respectively. Considering single-omics models, on average BLUP models (GBLUP and TBLUP) yielded less biased predictions than GBM models (GGBM and TGBM). For models integrating SNP genotypes and gene transcripts, GTBLUP and GTIBLUP showed similar bias across traits, while GTGBM had on average less bias than the BLUP models. For the GTCBLUP model, predictions were inflated (*β* < 1) for all traits but BMD21. In terms of prediction error, differences between models were smaller than observed for bias (Supplementary Table S2) or predictive accuracies (Table 4). The lowest RRMSE values were observed for FATP21, while the highest were observed for GLUC19. The RRMSE values for all traits analyzed were all around 1, indicating the average prediction errors were close to one phenotypic standard deviation.

#### Predictive ability for GEBV and other model components, and the correlation between them in BLUP models

In Table 5 the Pearson’s coefficient correlation between model components solutions (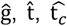 and ĝ_t_) for the different BLUP models and corrected phenotypes (**y**^*^) are shown. Overall, results for GTBLUP and GTIBLUP were similar across traits. These two models had the most accurate GEBV (ρ_ĝ_**y**_^*^) exclusively for GLUC8, while for BMD12 results from these models were matched by GTCBLUP. For GLUC19, all four parametric multi-omics models yielded the same accuracy for GEBV, which was the lowest (0.01) across traits. In 8 out of 13 traits the GEBV estimated using GTCBLUP model was the most accurate across all models. The correlation between 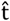 and 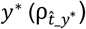 was also similar between GTBLUP and GTIBLUP, being always higher for these two models than observed for GTCBLUP. For GTCBLUP exclusively, 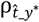 was low and negative for CHOL19 (−0.08), GLUC19 (−0.05) and TRGL8 (−0.07). For most traits, although a slight increase in the total variance explained was observed within the reference dataset (Figures 1 and 2) when comparing GTBLUP and GTIBLUP, there was not a proportional increase in ρ_ĝ_***y***_^*^ in the validation (Table 5). For GTCBLUP on the other hand, for all traits there was an increase in the variance explained by SNP genotypes (*g*^2^ in Figure 1 and Figure 2) when compared to GTBLUP and GTIBLUP, and the same pattern was observed for 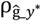. Results for 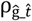 varied from +0.14 to +0.29 for GTBLUP and from +0.13 to +0.29 for GTIBLUP. For GTCBLUP, values for 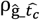 were all negative and close to zero, ranging from -0.13 to -0.03 (Table 5). The values for 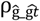, only calculated for GTIBLUP, were close to zero for most traits with an exception for CHOL8 and CHOL19, for which 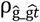 was 0.18. A similar pattern was observed for 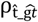, for which values varied from - 0.12 to +0.06, with the largest differences from 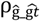 observed for CHOL8 and CHOL19.

**Table 5.**
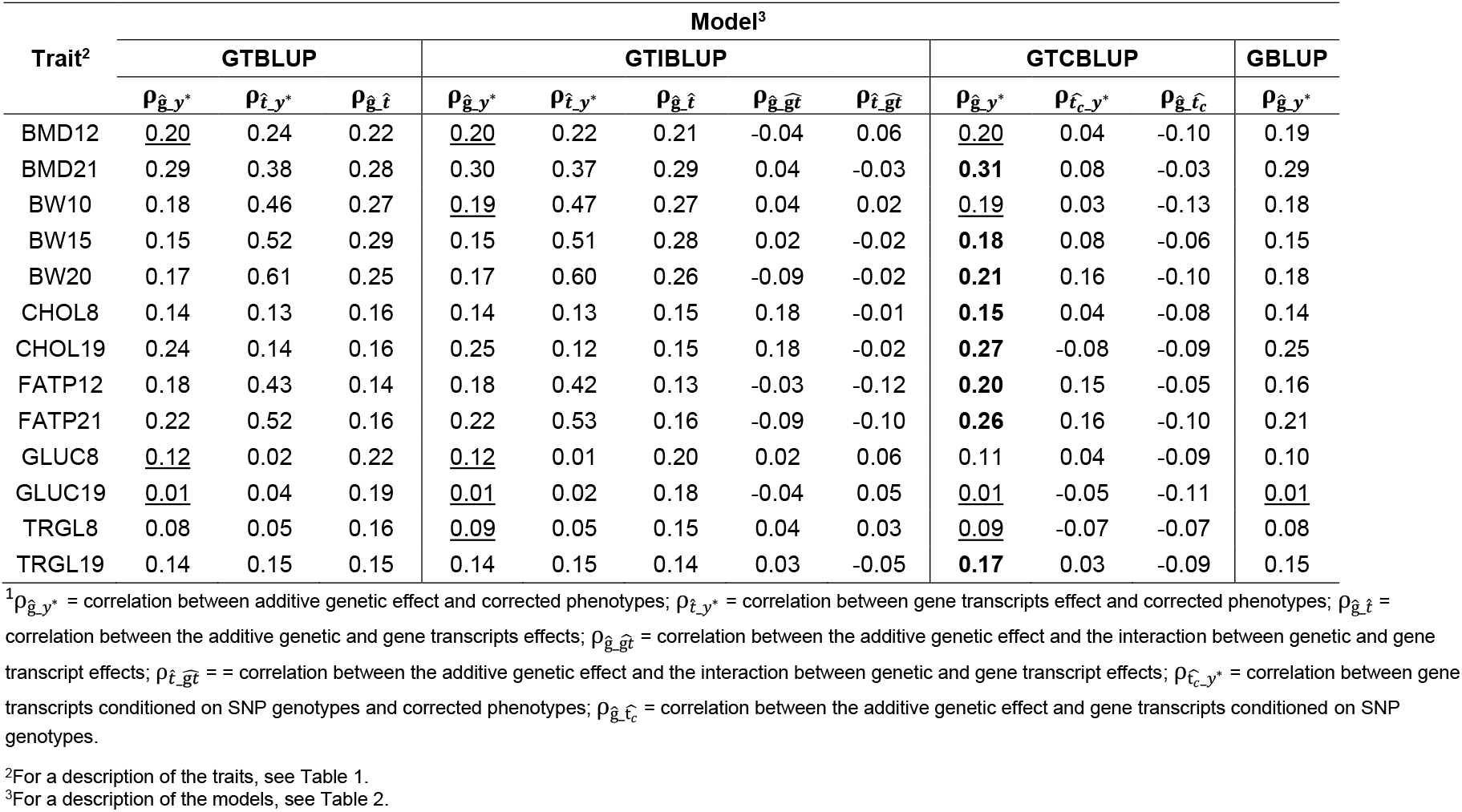
Pearson’s coefficient correlation (ρ) between model’s components 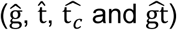 solutions and corrected phenotypes (**y**^*^) for BLUP models proposed^1^. Numbers in bold (per row) show the best values for the accuracy of GEBV 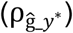 across models, identical results between two or more models are indicated in underline.

## DISCUSSION

Here, we investigated parametric and non-parametric approaches to leverage transcriptomic data for the prediction of complex phenotypes. To accomplish that, we used 478 animals from the DO Mouse population (Svenson *et al*. 2012), for which information on phenotypes (Churchill *et al*. 2012) for a wide range of quantitative traits, SNP genotypes and gene transcript levels from liver tissue (Tyler *et al*. 2017) were available on the same animals. We used the genomic (GBLUP) and transcriptomic (TBLUP) best linear unbiased prediction models to evaluate the value of these omics data to predict phenotypes. In addition, we evaluated models to integrate genome and transcriptome data by modelling both layers independently (GTBLUP) or including an interaction component between the genome and transcriptome (GTIBLUP). Finally, we proposed the GTCBLUP model that removes the between-omics-layer information redundancy. The gradient boosting machine (GBM) algorithm was investigated as a non-parametric approach potentially able to perform variable selection and capture non-linear effects by fitting either SNP genotypes (GGBM), gene transcript levels (TGBM) or to integrate both layers implicitly modeling interactions within and between omics layers (GTGBM).

Using data from six distinct traits measured at two or three time points (resulting in 13 traits in total), we first assessed the proportion of phenotypic variance explained by each variance component in the parametric models (Figure 1 and Figure 2). The variance explained by SNP genotypes and gene transcript levels (and their interaction) varied by trait, time of measurement and the model used. When using transcripts as predictors, two main patterns were observed. For 5 out 13 traits (Figure 1), the TBLUP model explained much more of the phenotypic variance than GBLUP. For the other 8 out of 13 traits (Figure 2), TBLUP explained less variance than GBLUP. The observation that the portion of variance explained by gene transcripts is strongly trait-specific is in line with results observed when assessing the proportion of variance from gene transcripts for complex traits in *Drosophila* (Morgante *et al*. 2020). Ehsani *et al*. (2012) have analyzed data from an F2 mice population using models integrating genotype markers and liver transcriptomics data. The authors reported that transcripts explained 79%, while genotypes explained 36% of the phenotypic variance for body weight at 8 weeks of age. It is important to emphasize that in Ehsani *et al*., (2012), RNA samples were measured at the same time point as phenotypes were collected. Other authors have observed that the genetic markers always explained a bigger portion of variance than transcripts in maize (Guo *et al*. 2016; Azodi *et al*. 2019) and *Drosophila* (Li *et al*. 2019).

There are several possible reasons for the conflicting results found in literature when assessing the value of transcripts to explain variation in complex phenotypes. Differently from genotypes, transcripts are affected by many factors such as the tissue from where samples are collected, the moment in life of sampling and the environmental conditions that the animal was exposed to. These variables most likely impact the variance explained (and concomitantly prediction performance) by transcripts. In Azodi *et al*. (2019) for example, transcripts were quantified from whole seedlings while phenotypes were recorded at a much older age, which could explain the limited predictive ability of transcriptomics data. In the present study transcripts were measured when the mice were 26 weeks of age, while all phenotypes were recorded at younger ages (from 8 to 21 weeks of age). In our results, phenotypes recorded closer to 26 weeks of age had a larger proportion of phenotypic variance explained by transcripts than measurements made earlier in the animal’s life for the same phenotype (Figure 1 and Figure 2). For BW and FATP the transcripts explained a larger proportion of phenotypic variance at all time points. For BMD, CHOL, GLUC and TRGL this was the case when there was 4 (BMD) to 6 weeks (CHOL, GLUC and TRGL) in-between measuring phenotypes and transcripts, while genomics explained more phenotypic variance when this time frame increased to 14 (BMD) or 18 weeks (CHOL, GLUC and TRGL) in-between. It is important to emphasize here that although this outcome may have been expected beforehand, to our knowledge it is the first time that this link between amount of variance explained by transcript versus the time difference between measuring transcripts and phenotypes has been shown empirically. One other aspect that must be considered here is that the gene expression from whole maize seedlings (Azodi *et al*. 2019) is probably much less related to traits collected later in life than the gene expression from liver tissue available for the DO mouse dataset. It is widely known that the liver is strongly linked to many metabolic pathways (Ponsuksili *et al*. 2019), and therefore likely also especially to the BW and FATP traits used here, while the variation contained in a sample collected from whole seedlings do not reflect a specific tissue but a pool of all tissues in this organism.

When fitting both SNP genotypes and gene transcripts as predictors the portion of variance explained by SNP genotypes varied drastically from the GBLUP model. For all BW and FATP traits, the proportion of variance explained by genotypes using GTBLUP and GTIBLUP was much lower than for GBLUP. Ehsani *et al*. (2012) and Takagi *et al*. (2014) observed a reduction in captured genetic variance by SNP genotypes of around 50% when fitting genotypes together with transcripts compared to models using fitting only genotypes as predictors for complex traits in other mice populations. This seems to confirm the hypothesis that there is redundant information between the genome and transcriptome layers (Wade *et al*. 2021), as also shown to be the case in *Drosophila* (Morgante *et al*. 2020). In our experience, it seems that the closer the phenotype analyzed is to the moment of RNA sampling, the higher the decrease in genetic variance captured by SNP genotypes in GTBLUP and GTIBLUP. This was observed for almost all traits we analyzed in different magnitudes. Takagi *et al*. (2014) analyzed circulating cholesterol at 10 weeks of age in mice and reported a large decrease in the genetic variance captured from SNP genotypes from models including only SNP genotypes (*g*^2^= 46%) and together with liver transcripts (*g*^2^= 19%) also measured at 10 weeks of age. In the present study, we observed only a slight decrease in genetic variance estimated when comparing GTBLUP (*g*^2^= 28%) and GBLUP (*g*^2^= 38%) for CHOL8. This seems to confirm that for the same phenotype, measurements made closer to the RNA sampling are prone to exhibit this pattern in a higher magnitude than measurements. This was further substantiated by the results observed for GTCBLUP. By conditioning the transcripts on the genotypes, the portion of variance explained by SNP genotypes was similar to the GBLUP model, while the variance explained by gene transcripts was much lower than estimated with TBLUP, GTBLUP and GTIBLUP.

The formal variance partitioning achieved with the BLUP models cannot be achieved with the non-parametric GBM models. To compare the performance of GBM and BLUP in terms of explained variance we investigated the model R^2^ within the reference set. For the GBM models it was almost always higher than for the BLUP models (Table 3). From our results, this pattern is recognizable for almost all traits analyzed, in which the GBM algorithm is able to capture a higher portion of variance than the parametric counterpart within the reference dataset (Table 3) but fails to outperform these models when predicting in the validation set (Table 4). The only exception to that is observed for GLUC19, for which in addition to the much larger portion of variance explained within the reference set, the GBM algorithm also outperformed BLUP models by a large margin for prediction purposes. The presence of noise in the data, limited size of the training set and the underlying complexity of the event being modelled are often cited as common causes of overfitting in machine learning models (Vabalas *et al*. 2019; Ying 2019). Here we used a training dataset of 286 animals and the high number of predictors in the models, coupled with the unavoidable presence of collinearity within and between omics layers may have caused GBM models to overfit. Takagi *et al*. (2014) have also analyzed datasets of similar size from another heterogeneous mice population using parametric models that integrated genomics and transcriptomics data, reporting large portions of phenotypic variance captured within training sets that weren’t necessarily translated to high predictive accuracy in the validation set for circulating glucose and cholesterol. The authors argue that the high number of model’s parameters to be estimated and the small number of animals with observations available could be the main cause of this pattern. We have observed a big impact of hyperparameters on the in predictive accuracies of the GBM models (results not shown). Having access to larger datasets could help to elucidate the magnitude of this impact for the models analyzed here since it would decrease the impact of hyperparameter definition in predictive performance, improving strength of evidence for any differences found between GBM and other models tested. The forward-prediction validation method adopted in the present study was previously described in Perez *et al*. (2022) and mimics prediction in animal and plant breeding, where a predictive model is trained based on data from individuals that have genomics (here also transcriptomics) and phenotype data available, while prediction is intended for un-phenotyped younger individuals. Here, training and validation generations were also 3 generations apart from each other, which erodes strong relationships (e.g., parent-progeny) and therefore should not be the cause of model overfitting suggested for the GBM models.

Prediction performance for phenotypes can be improved by combining genotypes and transcripts, however our results suggested that the magnitude of improvement is dependent on the trait analyzed (Table 4). In line with the observed differences in variance explained by model components, TBLUP showed a better predictive ability than GBLUP for most traits except CHOL19, GLUC8 and GLUC19. In contrast, Takagi *et al*. (2014) reported higher predictive accuracies for circulating glucose and cholesterol when using liver transcripts as predictors when compared to using genotypes in a different heterogeneous mice population (Valdar *et al*. 2006). Two aspects may explain the differences observed between studies. First, Takagi *et al*. (2014) performed a cross-validation scheme by randomly sampling individuals as reference and validations sets while we performed a forward-validation scheme, in which phenotype prediction for younger animals was based on estimates from older generations. Even though animals are randomly sampled, the overall similarity between reference and validation sets is higher in Takagi *et al*. (2014), which in general leads to higher magnitude of predictive accuracy (Pszczola *et al*. 2012; Werner *et al*. 2020). It is important to emphasize that although this is true when dealing exclusively with SNP genotypes, we cannot confirm that the affirmative holds when dealing with transcripts as predictors. A second relevant aspect is that in the present study phenotypes for CHOL and GLUC were collected at 8 and 19 weeks, while RNA samples were taken at 26 weeks of age. As previously mentioned, in Takagi *et al*. (2014) RNA samples and phenotypes were collected at the same age (10 weeks). In the present study, phenotypes recorded at a closer time point to transcript profiling result in higher predictive performance from transcripts (Table 4).

The TGBM model was able to overcome the TBLUP model for several traits, which was not the case when comparing GBLUP and GGBM. This result may indicate that interactions between transcripts were more easily captured or are more relevant than between SNPs. When fitting only genotype markers, the model is limited by incomplete linkage disequilibrium between the SNP and quantitative trait loci to perform an accurate detection of possible interactions, while there is no such limiting factor when using gene transcripts. One other hypothesis is that as transcripts are more strongly linked to phenotypes than genetic markers, transcript-by-transcript interactions are also likely to affect the phenotype more strongly than SNP-by-SNP interactions (Green *et al*. 2019), hence the former is expected to have a clearer and more detectable signal. Morgante *et al*. (2020) have used the random forest model, a non-parametric ensemble machine learning method like GBM, to predict complex phenotypes in *Drosophila* using gene transcripts as predictors but did not observe a superior predictive ability when compared to the TBLUP model. While TGBM consistently outperformed TBLUP in our study, the GTGBM was only partly outperforming GTBLUP and GTIBLUP. This could mean that the inclusion of SNP genotypes together with gene transcripts as predictors in the GTGBM model may have impaired the ability of GBM to capture linear and non-linear signals from within- and between-omics layers. The exact cause remains unclear, but the size of dataset together with the substantial increase in number of predictors when going from TGBM to GTGBM may be in the roots of it. It is likely that the GBM algorithm may require more data to be able to accurately capture all patterns from the complex relationship between omics layers underlying quantitative traits. If this is indeed the case, testing these models using a larger dataset could help to confirm this hypothesis. In Azodi *et al*. (2019) machine learning models integrating genomics and transcriptomics data were also not able to outperform single-omics models in terms of predictive accuracy for three traits in maize using a dataset of similarly limited size as in the present study.

The linear association for solutions from BLUP models’ components predicted for animals within the validation set were very similar between GTBLUP and GTIBLUP. For these two models 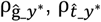 and 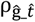 were almost identical across all traits (Table 5). Although the inclusion of an interaction component in GTIBLUP captured between 9% and 26% of phenotypic variance (represented by *gt*^2^ in Figure 1 and Figure 2) within the reference set, it did not seem to affect the relationship between other components. The low values observed for 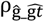 and 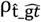 in GTIBLUP also seem to suggest that the interaction component is capturing a portion of variance not directly shared with ĝ or 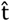 components, and therefore it does not affect the relationship between other components. The linear association between ĝ and 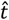 in GTBLUP and GTIBLUP models was always higher than the association between ĝ and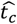 in GTCBLUP across all traits analyzed. Since in the GTCBLUP model the transcript relationship matrix was conditioned on the variance of SNP genotypes (*t*_*c*_), the linear association between solutions for the two components was expected to closer to zero than the observed in GTBLUP or GTIBLUP.

In this paper, we proposed the GTCBLUP model as an alternative to integrate genome and transcriptome data for genomic prediction. There has been an increasing interest in the use of intermediate omics data in animal and plant breeding (Guo *et al*. 2016; Yang *et al*. 2017; Azodi *et al*. 2019; Morgante *et al*. 2020; Christensen *et al*. 2021; Michel *et al*. 2021), such as transcriptomics, metabolomics, or microbiome data. The inclusion of new layers of omics data into genomic prediction models could arguably help in capturing additional portions of variance not explained by genotype data, but at the same time, these layers most likely contain overlapping information, increasing collinearity between predictors. Modelling the relationship between G and T components could be an efficient way to realize the added value of integrating such omics data into genomic prediction models (Wade *et al*. 2021), but this could also be a challenge given the increase in number of parameters to be estimated. The advantage of the GTCBLUP is that as pre-processing step it conditions the variance contained in transcripts on the variance of genotypes to minimize the amount of redundant information without having to increase model complexity. In general, the GTCBLUP model was able to produce GEBV that were at least as accurate as or slightly more accurate than the GBLUP model. The percentage of variance explained by SNP genotypes in GTCBLUP was similar to that with the GBLUP model, while it was always lower when using GTBLUP and GTIBLUP (Figure 1 and Figure 2). The observed reduction in additive genetic variance for GTBLUP and GTIBLUP when compared to GBLUP indicates strong redundancy in information contained in the genomic and transcriptomic layers. So, the conditioning of transcripts on SNP genotypes in GTCBLUP allowed this model to perform a more accurate variance partitioning for the additive genetic component, which consequently resulted in a more accurate estimation of GEBV (Table 5). An interesting alternative way to consider the covariance between genomics and transcriptomics layers, is by explicitly modelling it, as can be done using the CORE-GREML method (Zhou *et al*. 2020), implemented in the MTG2 software (Lee and van der Werf 2016). We considered this method to evaluate its potential, as well as to try and assess the magnitude of overlapping information between genotype and transcript data for the traits analysed. Results (Supplementary Table S3) indicated correlations from -0.47 to +0.71, with highest values observed for BW10, BW15, BW20, CHOL8 and CHOL19, FATP19 but the correlation coefficient was never significantly different from 0. Coincidently for most of these traits GTCBLUP obtained the most accurate GEBV when compared to GBLUP, GTBLUP and GTIBLUP. These trends seem to further support that, for specific traits, genomics and transcriptomics layers contain largely overlapping information, and although removing this redundancy does not result in more accurate phenotype prediction, it may contribute to obtain more accurate GEBV and consequently could improve breeding decisions. The lack of significance of the estimated correlations between genomics and transcriptomics data may be due to the limited size of the data used here, while based on the results obtained in the present study this does not provide a limitation for the GTCBLUP model.

One limitation of the GTCBLUP model is that it does not accommodate missing omics information, so all reference individuals must have genomics and transcriptomics data available. In the context of breeding programs, a situation in which all reference animals have multiple omics data available is unlikely to happen due to high costs involved in the collecting this kind of information. However, based on the observed decrease in costs of genotyping which has enabled large-scale genotyping, we may expect similar developments for the costs of transcriptomics and other intermediate phenotypes in the near future (Uzbas *et al*. 2019). At the same time, there have been some recent model developments that enable including other omics data in genomic prediction, when these other omics data are not available for all animals (Christensen *et al*. 2021; Zhao *et al*. 2022).

## CONCLUSION

We have assessed prediction models that incorporate genetic markers and transcriptomics data in genomic prediction of complex phenotypes in mice. The proportion of phenotypic variance explained by transcripts was almost always higher when traits were measured closer to the time of measuring gene transcripts. While GBM models explained more variance in the reference data, their predictive performance did not exceed the GBLUP models. Models including SNP genotypes and gene transcripts did not consistently outperform the best single-omics models to predict phenotypes. While TGBM model was able to outperform TBLUP, this was not the case for GTGBM compared to GTBLUP and GTIBLUP. The newly developed GTCBLUP model was able to force all phenotypic variance associated with SNP genotypes into its additive genetic component, by conditioning gene transcripts on SNP genotypes. GTCBLUP generally yielded considerably lower accuracies of phenotypic predictions than the other models including SNP genotypes and gene transcripts, but it showed the best accuracies for breeding values for most traits. We recommend using the GTBLUP model for prediction of phenotypes, while the GTCBLUP should be preferred when the aim is to estimate breeding values.

## SUPPLEMENTARY MATERIAL

**Table S1.** Regression coefficient of the corrected phenotypes on prediction for validation animals from all models tested. Values closer to 1 indicate less bias.

| Trait<sup>1</sup> | Model<sup>2</sup> |  |  |  |  |  |  |  |
| --- | --- | --- | --- | --- | --- | --- | --- | --- |
|  | Only SNP<sup>2</sup> |  | Only gene transcripts<sup>2</sup> |  | SNP + gene transcripts<sup>2</sup> |  |  |  |
|  | GBLUP | GGBM | TBLUP | TGBM | GTBLUP | GTIBLUP | GTCBLUP | GTGBM |
| BMD12 | 1.16 | 1.48 | 1.52 | 1.36 | 1.47 | 1.63 | 0.88 | 1.23 |
| BMD21 | 1.47 | 1.82 | 1.67 | 1.34 | 1.65 | 1.95 | 1.13 | 1.21 |
| BW10 | 1.07 | 1.46 | 1.25 | 1.46 | 1.28 | 1.44 | 0.85 | 1.16 |
| BW15 | 0.80 | 1.82 | 1.08 | 1.44 | 1.06 | 1.17 | 0.70 | 1.12 |
| BW20 | 1.01 | 1.82 | 1.09 | 1.36 | 1.09 | 1.16 | 0.82 | 1.07 |
| CHOL8 | 1.17 | 1.78 | 0.92 | 1.31 | 1.05 | 1.06 | 0.76 | 0.79 |
| CHOL19 | 0.99 | 0.79 | 0.79 | 1.43 | 0.92 | 1.00 | 0.83 | 0.69 |
| FATP12 | 0.55 | 0.76 | 0.96 | 1.16 | 0.99 | 1.09 | 0.89 | 1.18 |
| FATP21 | 1.10 | 0.83 | 1.02 | 1.25 | 1.04 | 1.12 | 0.99 | 1.14 |
| GLUC8 | 0.38 | 0.44 | 0.29 | 1.27 | 0.31 | 0.31 | 0.68 | 0.52 |
| GLUC19 | 0.35 | 1.82 | 0.33 | 1.28 | 0.36 | 0.37 | 0.26 | 0.45 |
| TRGL8 | 0.54 | 0.61 | 0.36 | 1.83 | 0.37 | 0.36 | 0.58 | 0.86 |
| TRGL19 | 0.96 | 1.12 | 0.75 | 1.15 | 0.74 | 0.85 | 0.72 | 1.23 |
| **Mean** | 0.89 | 1.27 | 0.93 | 1.36 | 0.95 | 1.04 | 0.79 | 0.97 |
| **Min** | 0.35 | 0.44 | 0.29 | 1.15 | 0.31 | 0.31 | 0.26 | 0.45 |
| **Max** | 1.47 | 1.82 | 1.67 | 1.83 | 1.65 | 1.95 | 1.13 | 1.23 |

<sup>1</sup>For a description of the traits, see Table 1.
<sup>2</sup>For a description of the models, see Table 2.

**Table S2.** Relative root-mean squared error (RRMSE) for predictions on the validation set. Lower values indicate lower prediction error.

| Trait<sup>1</sup> | Model<sup>2</sup> |  |  |  |  |  |  |  |
| --- | --- | --- | --- | --- | --- | --- | --- | --- |
|  | Only SNP |  | Only gene transcripts |  | SNP + gene transcripts |  |  |  |
|  | GBLUP | GGBM | TBLUP | TGBM | GTBLUP | GTIBLUP | GTCBLUP | GTGBM |
| BMD12 | 1.00 | 1.02 | 0.99 | 1.02 | 0.98 | 0.99 | 1.01 | 0.99 |
| BMD21 | 0.99 | 1.02 | 0.96 | 0.99 | 0.95 | 0.96 | 0.98 | 0.96 |
| BW10 | 0.88 | 0.90 | 0.85 | 0.87 | 0.85 | 0.85 | 0.88 | 0.85 |
| BW15 | 0.86 | 0.86 | 0.85 | 0.85 | 0.86 | 0.86 | 0.87 | 0.86 |
| BW20 | 0.87 | 0.88 | 0.84 | 0.86 | 0.84 | 0.84 | 0.87 | 0.84 |
| CHOL8 | 0.89 | 0.90 | 0.89 | 0.91 | 0.89 | 0.89 | 0.90 | 0.87 |
| CHOL19 | 0.95 | 0.97 | 0.96 | 0.97 | 0.95 | 0.95 | 0.96 | 0.95 |
| FATP12 | 0.91 | 0.91 | 0.86 | 0.87 | 0.86 | 0.86 | 0.89 | 0.87 |
| FATP21 | 0.85 | 0.87 | 0.77 | 0.78 | 0.77 | 0.77 | 0.81 | 0.77 |
| GLUC8 | 1.15 | 1.15 | 1.17 | 1.14 | 1.15 | 1.15 | 1.15 | 1.15 |
| GLUC19 | 1.31 | 1.27 | 1.33 | 1.27 | 1.31 | 1.31 | 1.34 | 1.31 |
| TRGL8 | 1.07 | 1.09 | 1.14 | 1.14 | 1.11 | 1.10 | 1.13 | 1.10 |
| TRGL19 | 1.19 | 1.22 | 1.18 | 1.16 | 1.18 | 1.17 | 1.20 | 1.21 |
| **Mean** | 0.99 | 1.00 | 0.98 | 0.99 | 0.98 | 0.98 | 1.00 | 0.98 |
| **Min** | 0.85 | 0.86 | 0.77 | 0.78 | 0.77 | 0.77 | 0.81 | 0.77 |
| **Max** | 1.31 | 1.27 | 1.33 | 1.27 | 1.31 | 1.31 | 1.34 | 1.31 |

<sup>1</sup>For a description of the traits, see Table 1.
<sup>2</sup>For a description of the models, see Table 2.

**Table S3.** Estimates of model parameters for the CORE-GREML model: the percentage of variance explained by SNP genotypes (*g*^**2**^) and gene transcripts (*t*^**2**^), the estimated correlation between g and t components (*ρ*_*gt*_), p-values for the estimated correlation (*p*[*ρ*_*gt*_]) and p-values for the likelihood ratio tests to determine whether the model fit by CORE-GREML was better than that by the standard model not including a covariance component (*p*[*LKH*]).

| Trait<sup>1</sup> | CORE-GREML parameters |  |  |  | GREML vs CORE-GREML |  |  |
| --- | --- | --- | --- | --- | --- | --- | --- |
| | $g^2$ | $t^2$ | $\rho_{gt}$ | $p[\rho_{gt}]$ | GREML | CORE-GREML | $p[LKH]$ |
| BMD12 | 0.24 | 0.20 | +0.21 | 0.62 | -181.4712 | -181.4062 | 0.71 |
| BMD21 | 0.28 | 0.26 | +0.14 | 0.55 | -177.2119 | -177.1537 | 0.60 |
| BW10 | 0.04 | 0.40 | +0.58 | 0.29 | -695.5398 | -695.3162 | 0.50 |
| BW15 | 0.03 | 0.45 | +0.60 | 0.45 | -696.1394 | -696.7851 | 0.25 |
| BW20 | 0.01 | 0.60 | +0.63 | 0.32 | -679.8870 | -680.7453 | 0.19 |
| CHOL8 | 0.27 | 0.13 | +0.55 | 0.45 | -483.6665 | -483.9623 | 0.44 |
| CHOL19 | 0.21 | 0.18 | +0.71 | 0.07 | -388.3944 | -389.8146 | 0.09 |
| FATP12 | 0.17 | 0.73 | -0.23 | 0.50 | -17.1618 | -17.0426 | 0.62 |
| FATP21 | 0.08 | 0.89 | -0.47 | 0.34 | -3.5900 | -3.4460 | 0.59 |
| GLUC8 | 0.21 | 0.09 | -0.22 | 0.76 | -510.4316 | -510.4436 | 0.87 |
| GLUC19 | 0.02 | 0.15 | -0.26 | 0.72 | -402.9910 | -403.0818 | 0.67 |
| TRGL8 | 0.08 | 0.14 | +0.26 | 0.23 | -226.2193 | -226.9104 | 0.25 |
| TRGL19 | 0.09 | 0.25 | +0.07 | 0.90 | -171.9591 | -171.9595 | 0.97 |

<sup>1</sup>For a description of the traits, see Table 1.

